# Controlled “out-of-season” spawning of reef-forming corals in aquaria using offset environmental profiles

**DOI:** 10.1101/2024.12.05.626918

**Authors:** Lonidas P. Koukoumaftsis, Matthew Salmon, Glenn Everson, David J. Hughes, Deepa Varkey, Andrew J. Heyward, Andrea Severati, Craig Humphrey

## Abstract

The global climate crisis has heightened the urgency for developing interventions to enhance resilience and recovery of coral reef ecosystems. However, research programs are often bottlenecked by availability of coral early life stage material due to the seasonal (annual) nature of coral mass spawning events. Here, we present a proof-of-concept of “out-of-season” coral spawning, utilising advanced aquarium control technology to induce spawning in multiple key Great Barrier Reef (GBR) coral species held in long-term indoor aquaria. By developing a 6-month offset environmental profile encompassing seasonal temperature, photoperiod, and lunar profiles, we successfully induced synchronised mass coral spawning during austral autumn/winter (May - June) over consecutive years in 2022 and 2023. We also “phase-shifted” the hour of sunset by five hours on spawning nights, creating a more favourable time window (i.e., minimising late nights) for gamete fertilisation. The timing of gamete release in captive corals closely matched previous GBR field observations for both nights after full moon (NAFM) and time after sunset (TAS). Gamete fertilisation was successful for six GBR species: *Acropora millepora, Acropora loripes, Acropora hyacinthus, Acropora elseyi, Acropora austera,* and *Montipora aequituberculata* producing a total of 1.75 million larvae. We outline key physiological insights gained into environmental regulation of coral spawning synchronicity and discuss the potential for out-of-season spawning to accelerate various aspects of coral reproduction research and enhance the growing toolbox of active restoration strategies aimed at reversing global coral reef decline.

## Introduction

Coral reefs are hotspots of biodiversity with immense socio-economic value yet are facing increasing pressures globally due to climate change (Hoegh-Guldberg et al., 2017; Hughes et al., 2018, 2020) and localised impacts from coastal urbanisation (Rosenberg et al., 2022). According to the IPCC, up to 99% of reefs are expected to be degraded by 2100 even under optimistic climate scenarios (Masson-Delmotte et al., 2018). The coral reef crisis has placed urgency in efforts to document acute and chronic disturbances, understand the ecophysiological responses of corals to environmental change and develop interventions to facilitate resilience and recovery of reef ecosystems (Sweet and Brown, 2016; Randall et al., 2020; McLeod et al., 2022).

However, within populations our understanding of the physiological responses of coral larvae and recruits remains comparatively less well-developed than for their adult counterparts (e.g., McLachlan et al., 2020). This represents a major knowledge gap since the success of coral early life stages is critical in underpinning the long-term viability of reef assemblages at local scales (Hughes and Tanner, 2000). One of the major bottlenecks in improving our understanding of reef resilience and recovery is the limited opportunity to conduct coral spawning research owing to the ephemeral nature of these events. Most reef-building corals (Order: Scleractinia) are broadcast spawners, exhibiting synchronous gamete release within a population often involving multiple species spawning on the same few nights each year, commonly referred to as “mass spawning” events (Baird et al., 2021). Synchronisation of spawning events is a critical reproductive strategy to maximise cross-fertilisation opportunities between coral conspecifics and is thought to align with optimal environmental conditions for survival, dispersal and settlement of coral larvae (Oliver and Babcock, 1992; Miller and Mundy, 2003; Harrison, 2011).

The process of synchronous gamete release by corals appears to be finely regulated by a sequence of environmental cues that operate over timescales ranging from hourly to seasonal (Fogarty and Marhaver, 2019). For example, the month in which spawning occurs is most closely correlated to seasonal variability in sea surface temperature (SST), the specific day of spawning appears strongly linked to factors associated with the lunar cycle (e.g., tides and moonlight intensity), while the time-of-day is likely associated with the period of darkness immediately following sunset (Harrison et al., 1984; Babcock et al., 1986; Oliver and Babcock, 1992; Van Woesik et al., 2006; Fogarty and Marhaver, 2019). Regional variation in environmental cues has also been observed, with solar rhythms playing a more prominent role than SST in regulating spawning synchronisation among Caribbean corals, suggesting the influence of environmental driver(s) may differ by location, species, or both (Van Woesik et al., 2006).

Four decades on from the realisation that most coral species are broadcast spawners (Harrison et al., 1984; Babcock et al., 1986), access to spawning events underpins a broad range of coral research. Spawning events are generally annual and usefully predictable - albeit slightly variable each year for a given location (Baird et al., 2021) – however, the episodic nature of coral spawning nonetheless poses a challenge for marine scientists investigating coral reproduction and recruitment processes. While the timing of mass coral spawning events is well-documented on reefs in both hemispheres, the seasonality which is a feature of broadcast spawning at all latitudes, including equatorial reefs (e.g., Guest et al., 2002), limits the window of opportunity for researchers to access coral gametes. For example, if a coral species of interest has a single spawning period, researchers may have just one opportunity per year at that particular location. Seasonal constraint on gamete availability therefore imposes a bottleneck for research programs requiring access to coral early life history stages (Cleves et al., 2020), particularly where it is desirable to work with local species or genotypes (e.g., Quigley et al., 2020; Quigley and van Oppen, 2022).

Control over environmental parameters has been used to manipulate reproductive cycles of various marine organisms, notably those in aquaculture hatchery systems (Firkus et al., 2024). Advances in aquarium control technology, allowing for precise replication of natural environmental profiles, together with improved understanding of how environmental factors regulate coral spawning synchronicity, have enabled researchers and hobbyists alike to induce predictable spawning of aquarium corals (Craggs et al., 2017). To date, multiple Indo-Pacific coral species have been spawned in aquaria in this fashion, typically releasing gametes on the same night of the year as their natural reef counterparts in the wild and often within minutes of observed release times in nature (Baird et al., 2021; Sakai et al., 2024).

Ability to replicate and control seasonal variability in temperature and photoperiod (i.e., photothermal manipulation, Martin-Robichaud and Berlinsky, 2004) within aquaria allows for development of protocols to induce captive coral spawning at different times of the year to achieve “out-of-season” (or “off-season”) spawning. Out-of-season spawning of corals could be utilised to provide a controlled source of gametes and larvae at pre-determined times of the year providing an important boost for relevant research programs, yet this possibility has been poorly explored to date. We conducted an 8-year program of research culminating in the development and application of seasonally offset environmental profiles to trigger synchronised and predictable spawning of multiple Great Barrier Reef (GBR) coral species in aquaria during austral autumn/winter (May - June). This breakthrough addresses the issue of seasonality of larval supply by expanding the availability of coral larvae beyond their natural spawning window(s). We highlight how out-of-season spawning could provide an immediate boost to coral research programs reliant upon early life stage material and out-of-spawning systems, and open novel possibilities to gain deeper insight into various aspects of coral reproductive physiology.

## Materials and Methods

### Generation of broodstock

All experimental work was conducted at the National Sea Simulator (SeaSim), located at the Australian Institute of Marine Science (AIMS, Townsville, Australia). For broodstock, we selected first filial generation (F1) adult coral colonies that had been reared and maintained in captivity for ∼5 years and thus exhibited a strong track record of growth and survival within SeaSim. A total of six coral species were included in this research program: *Acropora hyacinthus*, *Acropora millepora*, *Acropora loripes*, *Acropora elseyi*, *Acropora austera* and *Montipora aequituberculata*. F1 corals were established from gametes obtained from parental (F0) corals collected from the central Great Barrier Reef at Davies Reef (mid-shelf; 18.820° S, 147.645° E), Backnumbers Reef (mid-shelf; 18.508° S, 147.148° E), and the Palm Islands region (inshore; 18.772° S, 146.532° E) between 2014-2018, 3-7 nights before the October, November, or December full moons (Great Barrier Reef Marine Park Authority [GBRMPA] permit G21/38062.1). On average, 12 parental colonies (F0) of each species were collected by SCUBA divers using hammer and chisel, transported to SeaSim, and held in 1200L outdoor aquaria maintained as described in Ramsby et al., (2024). During spawning periods each year between 2014-2018 (Fig. 1), after sunset, broodstock corals were monitored for visible formation of egg/sperm bundles under the polyp discs (i.e., “setting” *sensu* Harrison et al., 1984). Once setting was observed, colonies were isolated, bundles collected and fertilised as described in Heyward and Negri (1999).

**Figure 1:**
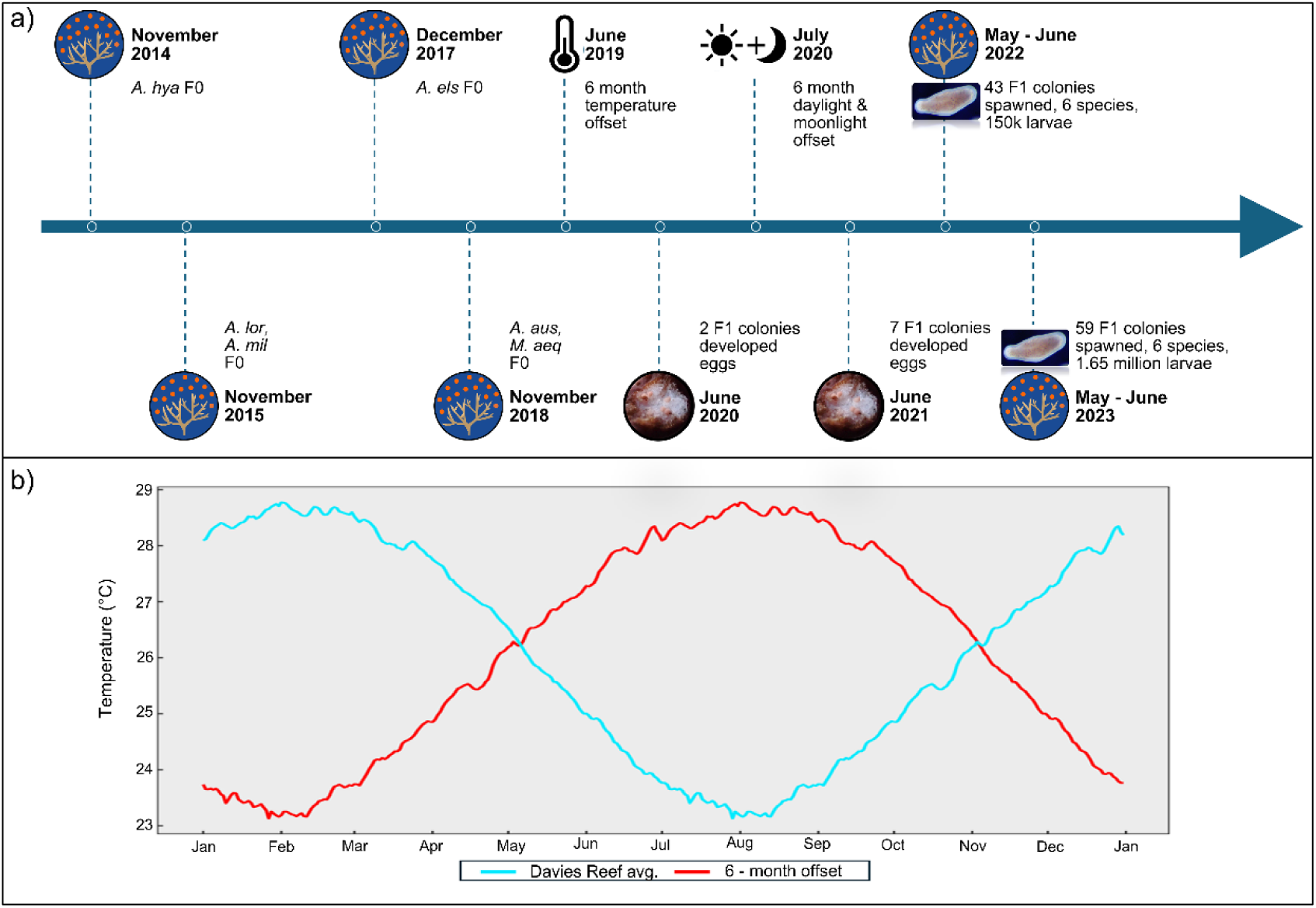
a) Timeline highlighting key events associated with the “out-of-season” spawning project conducted at the National Sea Simulator (SeaSim) between 2014-2023. b) Temperature profile for Davies Reef (Central Great Barrier Reef, Australia, -18.828 S, 147.643 E) consisting of data averaged between 1998 - 2015 (blue line) and the same dataset with a 6-month temporal offset applied to generate the out-of-season temperature profile (red line). Davies Reef temperature data was acquired from the Australian Institute of Marine Science (AIMS) Davies weather station, where measurements corresponded to a depth of 4m (https://weather.aims.gov.au/#/station/4).

Fertilisation was verified by visually inspecting a subsample (*n* = 100) of fertilised eggs (embryos) for early stages of cleavage (cell division) indicative of embryogenesis. Once fertilisation was established, embryos were skimmed from the water surface, washed, and transferred to 85 L culture tanks following handling procedures as described in Ramsby et al., (2024). At 48 h post-fertilisation, embryos had developed into free-swimming planula larvae, and approximately one-week post-fertilisation larvae were settled onto pre-conditioned aragonite plugs (20 mm diameter, Ocean Wonders). Aragonite plugs had been pre-conditioned for eight weeks in an outdoor aquaria system to achieve 30-60 % coverage of crustose coralline algae (CCA) - a known cue that induces the coral settling response (Heyward and Negri, 1999; Webster et al., 2004; Abdul Wahab et al., 2023). Settlement was performed in 50 L acrylic tanks housing PVC plug trays – each holding 165 aragonite plugs (Fig. S1) and seawater exchange was provided continuously at a low flow rate (0.2 L min^-1^). Approximately 1200 coral larvae were introduced into each 50 L acrylic tank, and larvae were deemed to be sufficiently attached to the aragonite plugs after 24 h, at which point the rate of seawater exchange was increased to 0.8 L min^-1^ (providing a complete system exchange per hour). After a further 48h, PVC plug trays containing settled larvae were transferred to indoor community, semi-closed aquariums (280 L, provided with three system turnovers per day of seawater). These community aquaria were equipped with a protein skimmer to remove organic waste and augment gas exchange (Hughes et al., 2022), while mechanical filtration was provided by a media filter system containing glass beads (∼0.7 µm diameter). To facilitate uptake of photosynthetic endosymbionts (Family: Symbiodiniaceae), fragments of F0 parental colonies were introduced as donor colonies alongside newly settled recruits (Williamson et al., 2021). Corals were maintained in community aquaria under a constant temperature regime (27.25 ± 0.3°C) for five years between 2014-2019 until being moved to a dedicated experimental holding room.

### Experimental setup – Out-of-season spawning

Adult F1 corals (See Fig. 2 for numbers/species) were moved to a dedicated experimental system with the capability to dynamically manipulate (and offset) key environmental variables including water temperature, photoperiod and moonlight. Corals were distributed across two aquarium systems, each comprising two 1200 L tanks connected to a common 700 L sump. Briefly, each system was supplied with 1 µm filtered seawater and maintained at a salinity of 35 ± 1 PSU. Biological filtration capacity was provided by ceramic biological media (MarinePure®, CerMedia, Buffalo, NY, USA). Seawater temperature was continuously monitored by a PT100-type FEP-insulated resistance temperature detector (Omega, Deckenpfronn, Germany) interfaced to a PCS7 Siemens Supervisory Control and Data Acquisition (SCADA) control system. Dynamic control of aquarium temperature was achieved using logic feedback applied to electric actuating valves, allowing a variable percentage of cold (15°C) or hot (40°C) water to pass through a shell-and-tube heat exchanger system (Fig. S2) to achieve system temperature within ±0.3°C of the programmed setpoint. System water was continuously recirculated by a centrifugal magnetic drive pump (MX 400, Iwaki, Tokyo, Japan), providing flow of 40 L min^-1^ to each tank, with ∼90 L min^-1^ passing through the heat exchanger (Fig. S2). In-tank water circulation was provided by two gyre pumps (XF280, Maxspect, Hong Kong, China). Seawater was continuously exchanged at a rate of 1.7 system turnovers per day to ensure stability of key water chemistry parameters essential to coral growth (e.g., alkalinity, calcium and magnesium, see Craggs et al., 2020; Hughes et al., 2022). Levels of nitrate (NO_3_^-^) and phosphate (PO_4_^3-^) were measured weekly using a segmented flow analyser (Seal AA3, Norderstedt, Germany) and did not exceed 5 µM and 0.5 µM respectively throughout the study except during periods of gamete release where in-tank nutrient levels were temporarily elevated (Fig. S3). Corals were provided with a range of feeds once a day including microalgae (comprising a mixture of *<u>Dunaliella</u>* sp., *Tisochrysis lutea,* and *Chaetoceros muelleri*) at a final density of 2000 cells mL^-1^), HUFA-enriched instar II *Artemia salina* (0.5 nauplii mL^-1^), unenriched instar I *A. salina* (0.5 nauplii mL^-1^) and phytoplankton-enriched rotifers (0.35 nauplii mL^-1^). Corals were held under a maximum photon irradiance (400-700 nm) of 150 µmol photons m^-2^ s^-1^ while photoperiod was adjusted weekly to match local conditions in Townsville, Australia (Time and Date AS 2022). One system was fitted with 16 commercially available LED panels (Hydra® 52, Aqua Illumination®, Ames, IA, USA), while the second was fitted with eight custom-designed LED panels (see Fig. S2 for a comparison of measured spectra). Initially, corals were held at a constant temperature of 27.25°C (i.e., matching that of the previous holding aquaria), until an offset (out-of-season) temperature profile was applied (Fig. 1).

**Figure 2:**
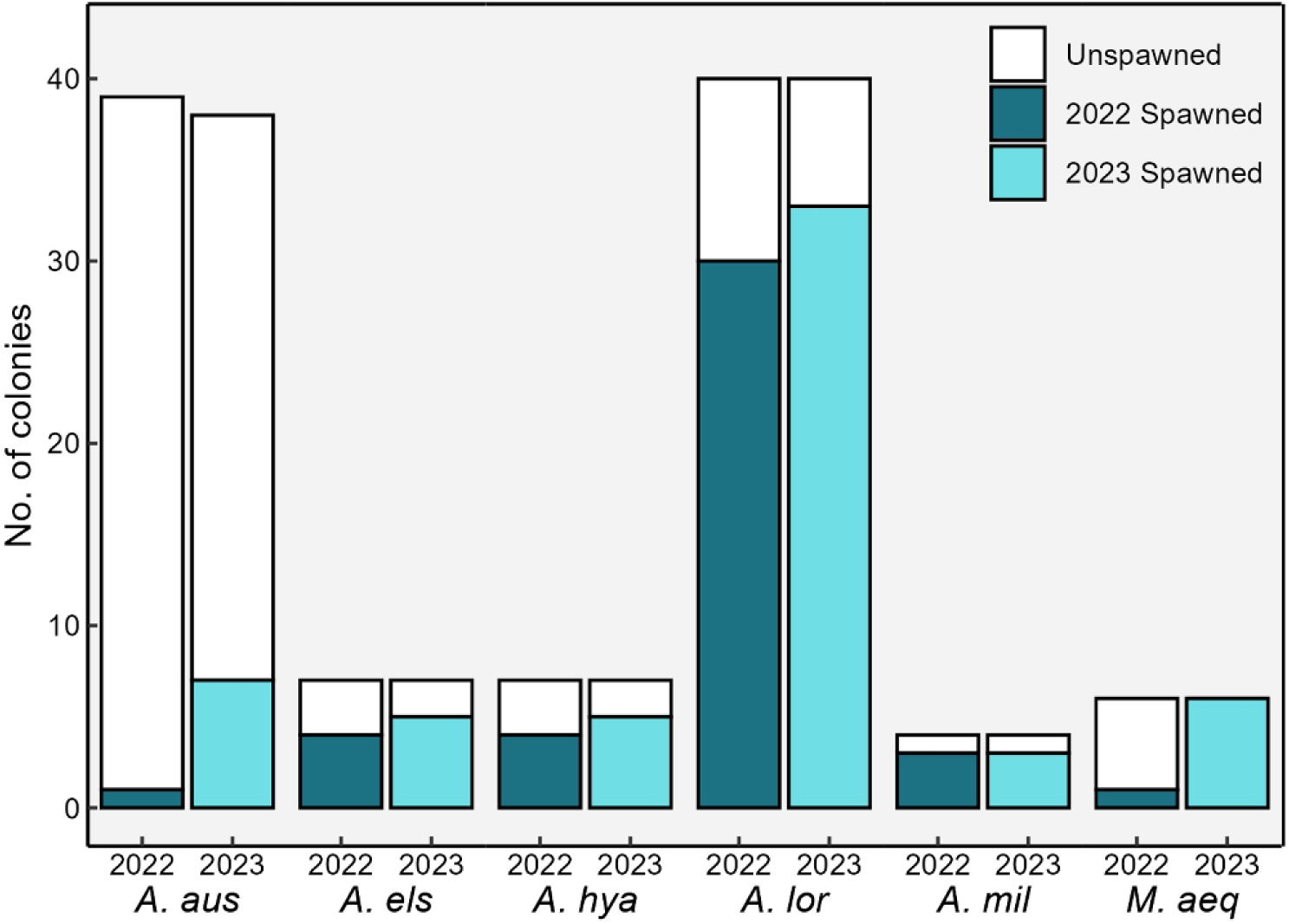
Total number of Filial 1 (F1) broodstock colonies in the out-of-season spawning project and proportions spawned in 2022 (dark green) and 2023 (turquoise). N.b. the number of *A. austera* colonies in 2023 (n = 38) was lower than in 2022 (n = 39) due to mortality. Species key*: A. aus* – *Acropora austera. A. els* – *Acropora elseyi. A. hya* – *Acropora hyacinthus. A. lor* – *Acropora loripes. A. mil* – *Acropora millepora. M. aeq* – *Montipora aequituburculata*.

### Applying an “out-of-season” offset temperature profile

To create a representative seasonal temperature profile, historical temperature data were sourced from the AIMS weather station located at Davies Reef (18.832° S, 147.635° E). Specifically, data from 4m depth over an 18-year period (1998-2015) were acquired and averaged to generate a composite seasonal temperature profile reflective of Davies Reef (representative of the central GBR region, Fig. 1). This composite profile was then uploaded as a lookup file to a programmable logic controller (PLC) permitting dynamic control of aquarium water temperature via SCADA as described above. To achieve an offset (i.e., out-of-season) profile, we advanced the date of the PLC by six months, thus daily lookup temperature values for experimental systems were six months ahead of the true date. In June 2019, the out-of-season (offset) temperature profile was initiated in both experimental systems (Fig. 1). This date was chosen because the temperature difference between the fixed temperature (27.25°C) and the out-of-season temperature profile (27.27°C) on this day of year was minimal (Fig. 1), thus minimising any physiological stress to the corals. Out-of-season temperature profiling was run for 13 months from June 2019 – July 2020, after which offset profiling of photoperiod and moonlight was also incorporated (Fig. 1).

### Offsetting photoperiod and moonlight

In July 2020, dynamic profiling of photoperiod and moonlight was introduced (Fig. 1). A six-month offset photoperiod profile was designed to match the 6-month future photoperiod at Townsville, Australia (Time and Date AS 2022). We also phase-shifted the photoperiod, bringing sunrise/sunset forward by four hours to provide a more convenient time window for researchers to observe spawning and harvest gametes for research.

To replicate the very low natural levels of moonlight experienced on the reef (Sweeney et al., 2011), we used a custom-made white LED fitted with a diffuser and positioned to face the ceiling, thus providing corals with scattered incident light. Moonlight intensity was calibrated to a maximum of 1 lux at full moon as measured in air using a portable lux sensor (3281A, Yokogawa, Tokyo, Japan). Profiling control of moonlight was achieved using a custom dimmable LED driver to provide a given percentage of 1 lux relative to the lunar phase for a full moon. A moonlight profile was created using predicted future moonlight data (https://www.timeanddate.com) loaded into the PLC to allow automatic profiling of moonrise/moonset times matching those of the six-month future moon in Townsville, Australia. Importantly, because the corals had been held under a fixed lighting regime over multiple years, we opted not to replicate full seasonality of insolation whereby photoperiod was adjusted to reflect seasonal variance in daylength, but daily irradiance maximum remained constant year-round.

### Verifying gametogenesis, F1 spawning, and F2 generation

In the week prior to expected spawning in May (2020-2023), two 3-4 cm nubbins per colony were removed from selected *A. loripes, A hyacinthus, A. elseyi*, *A. millepora, A. austera, and M. aequituberculata* colonies. Nubbins were fixed in a 4% formaldehyde-saltwater solution for 24 h and then decalcified over seven days in a 3% hydrochloric acid-saltwater solution. Branches from coral samples were subsequently cross-sectioned and oocyte development was examined under a stereo microscope (Leica MZ16A; Leica Microsystems GmbH, Wetzlar, Germany) (Fig. 3).

**Figure 3:**
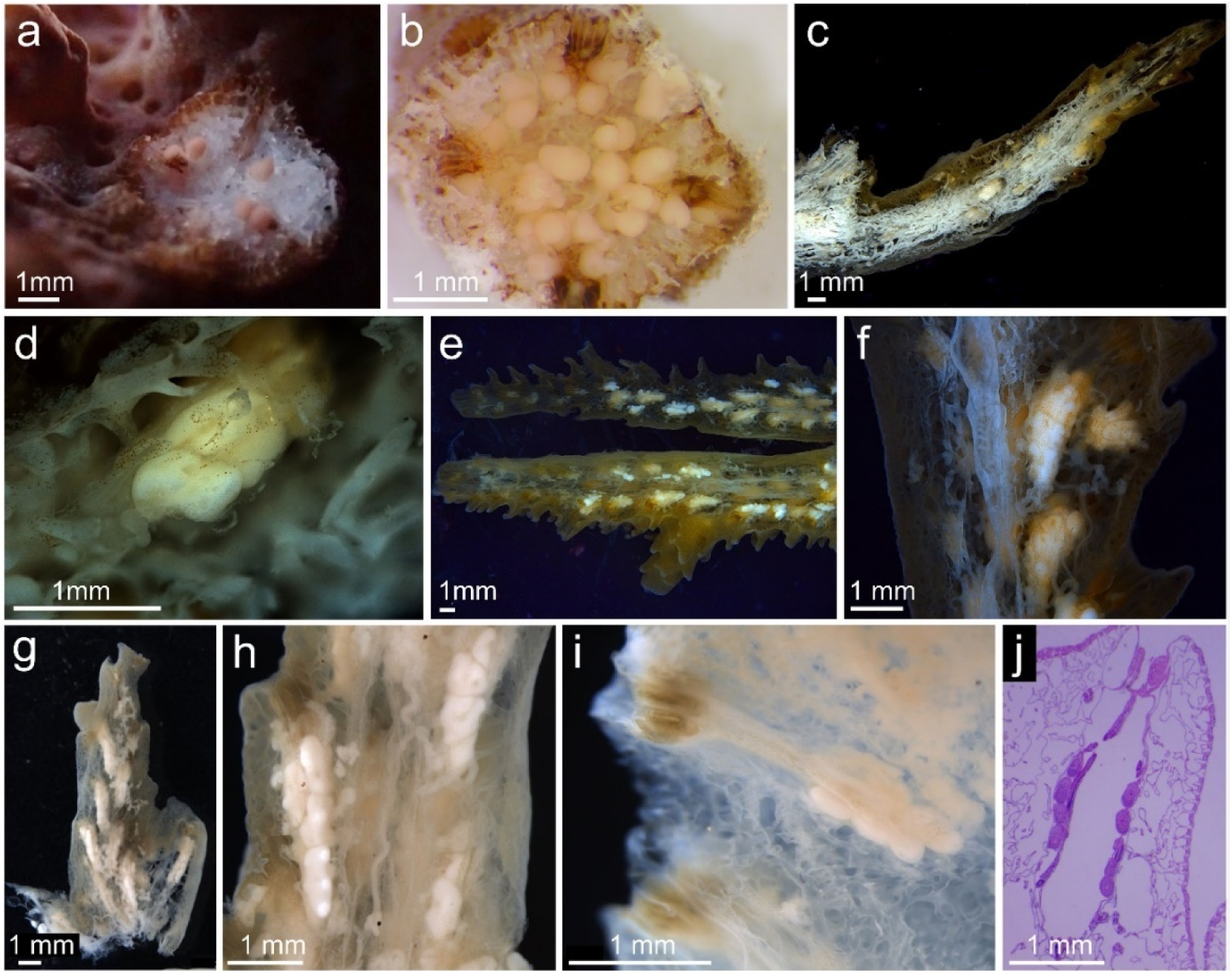
Out-of-season coral broodstock oocyte development showing (a) presence of pigmented eggs in *Acropora hyacinthus* (June 2020), (b) presence of pigmented eggs *in Acropora millepora* (June 2020), (c-d) decalcified *A. hyacinthus* samples showing developing eggs (June 2021), (e-f) decalcified *A. millepora* samples showing developing eggs (June 2020), (g-h) decalcified *A. hyacinthus* samples showing developing eggs (May 2021), (i) decalcified *A. loripes* sample showing developing eggs (May 2021), (j) decalcified and sectioned *A. loripes* samples prepared for histological analysis showing presence of developing eggs (April 2022).

During 2022 and 2023 spawning events (Fig. 1), F1 broodstock corals were observed, isolated (where “setting” was apparent), gametes collected, fertilisation attempted and verified as described above. Fertilisation success was determined as the proportion (%) of embryos relative to the total egg sample. Larval cultures were made as above, and a subset of second filial generation (F2) larvae were settled to achieve life cycle closure of captive coral species (see Supplementary Methods).

### Statistical analyses

To evaluate how spawning dates (expressed as nights after full moon, NAFM) and times (minutes after sunset) of corals in this study aligned with observations elsewhere, we compared them to data sourced from the Indo-Pacific coral spawning database (Baird et al., 2021). Specifically, we filtered observations from the GBR ecoregion for each species in our study, corresponding to the same geomorphological reef groups that our F0 broodstock originated from, i.e., inshore vs. mid-shelf reefs. An exception was made for *A. austera*, for which we included observations of both inshore and mid-shelf reefs due to the low overall number of observations within the database (*n* = 3 total, comprising two inshore and one mid-shelf observations). For *A. elseyi*, three observations of NAFM were excluded because we were unable to reconcile the reported date of the full moon with those listed in the original sources. To understand how closely spawning in our study aligned to previous observations, we calculated the difference (ΔNAFM) between NAFM values for each of our corals that spawned, against the average NAFM value reported in Baird et al., (2021) for the same coral species. A difference of one night (whether earlier or later than the Baird et al., (2021) average value) was scored as +1, thus always yielding a positive ΔNAFM value. To assess predictability of spawning in our out-of-season spawning program, ΔNAFM values were fitted to a Generalised Linear Model (GLM) assuming a Poisson distribution (with log link function) using R software (v4.3.1). Multiple comparisons with Bonferroni correction were performed on the adjusted least-square means using the emmeans software package (v1.8.8) to examine differences between groups of interaction terms.

## Results

### Spawning observations (by year)

#### 2020

No coral spawning was observed during 2020, however visual observations of pigmented eggs coupled with decalcified samples taken on 21^st^ June 2020 (21^st^ December equivalent) confirmed late-stage gametogenesis in a single colony of both *A. hyacinthus* (Fig. 3 a, c-d) and *A. millepora* (Fig. 3 b, e-f). Whether gametes were released from these gravid corals at an unexpected time (and thus were not observed) or were reabsorbed by the parental colonies (e.g., Okubo et al., 2007) was not resolved, but the former appears more likely given the rarity of gamete reabsorption in the absence of significant environmental or mechanical stress (Bauman et al., 2011).

#### 2021

As in 2020, no spawning was observed in 2021 despite the presence of developing eggs in two *A. hyacinthus* and five *A. loripes* samples (Fig. 3 g-i). As in the previous year, it was unclear if gamete release was not observed due to the corals spawning at an unexpected time or gametes were reabsorbed by the parental colonies. Importantly, it was later discovered that there had been an error in the moonlight data lookup table, where moonrise/moonset times were incorrect for any given day in the three months leading up to spawning, where critically, the full moon (1 lux intensity) cue was not delivered on the correct date.

#### 2022

A total of 43 captive coral colonies (out of 109) spawned in the out-of-season program during 2022 (Fig. 2, Table S1). Spawning was split across two months (May and June), with the majority (38 of 43) of colonies spawning in June. Between 13^th^-16^th^ May (6-9 NAFM) four coral species spawned, comprising three colonies of *A. elseyi*, two colonies of both *A. millepora* and *A. loripes*, and a single colony of *A. austera* (see Table S1). Notably, some corals partially spawned over multiple nights (e.g., one colony of *A. millepora* spawned over three successive nights) (Table S1). Fertilisation success ranged from 32% (*A. elseyi*) to 80% (*A. millepora)* (Table 1, see also Fig. 4, Table S2). Fertilisation was not attempted for *A. loripes* colonies as only a small number of bundles were released (i.e., a partial spawn), or for *A. austera* where only a single colony spawned. In total, 25,000 larvae were produced in May 2022 comprising 15,000 and 10,000 from *A. millepora* and *A. elseyi* respectively (Table 1).

**Figure 4:**
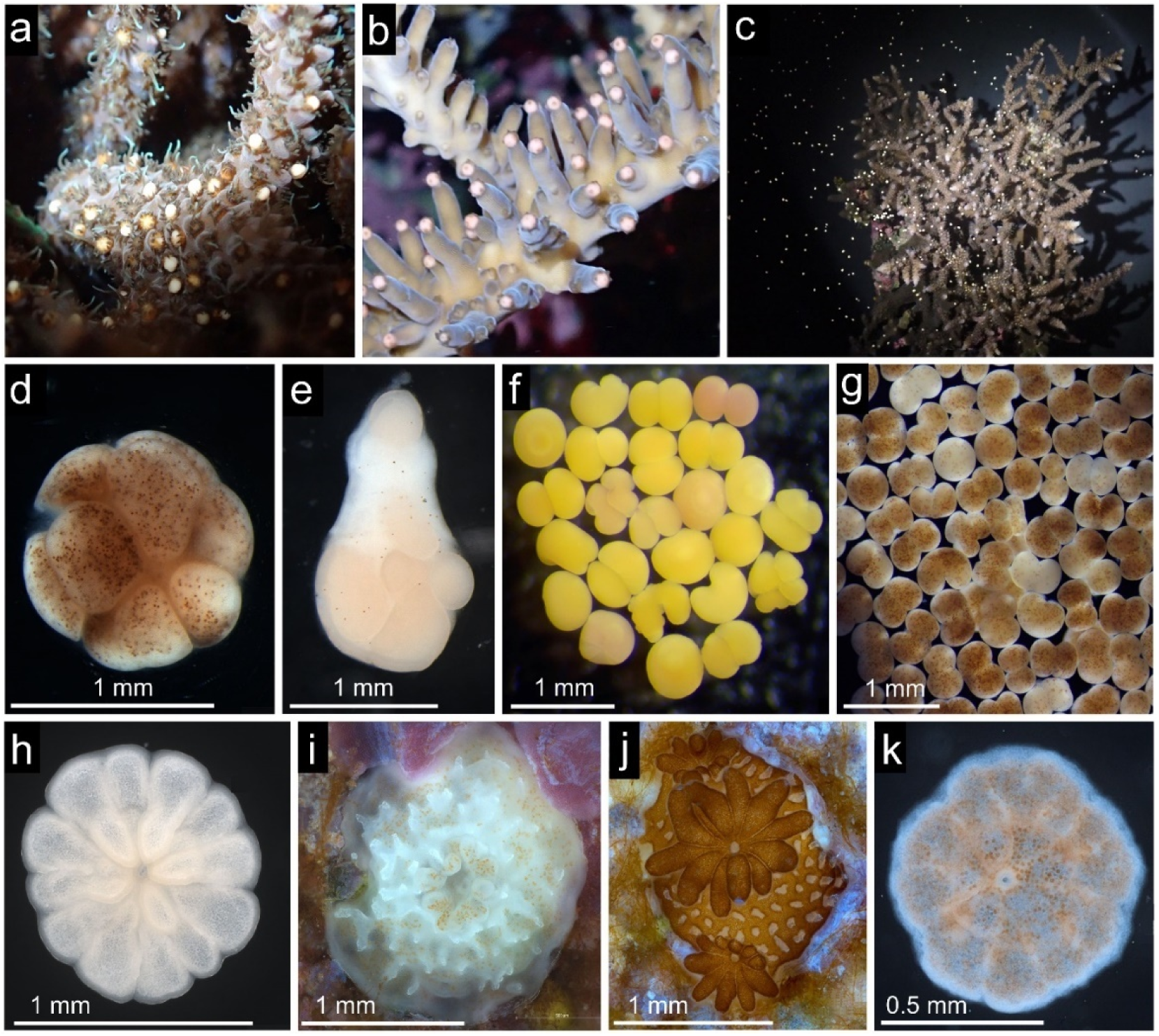
Out-of-season spawning, fertilisation, and filial 2 (F2) recruits (a) *Acropora millepora* adult colony “setting” (*sensu* Harrisson et al., 1984) (May 2022), (b) *Acropora elseyi* adult colony “setting” (May 2022), (c) Gamete release from *A. millepora* adult colony (May 2022), (d) Close up photograph of *Montipora aequituberculata* bundle (June 2022), (e) Dissociation of *Acropora loripes* bundle (June 2022), (f) *A. millepora* fertilised embryos (May 2022), (g) *M. aequituberculata* fertilised embryos (May 2023), (h) *A. millepora* F2 recruit (May 2022), (i) *Acropora hyacinthus* F2 recruit (June 2022), (j) Established *A. loripes* F2 recruit showing budding growth and established Symbiodiniaceae population (19 days post-settlement - June 2022), and (k) *M. aequituberculata* F2 recruit (May 2023).

**Table 1:** Summary of larval production from out-of-season spawning (2022-2023) showing the number of larval cultures established, fertilisation percentage range*, and total larvae numbers generated. Species key: *A. aus – Acropora austera. A. els – Acropora elseyi. A. hya – Acropora hyacinthus. A. lor – Acropora loripes. A. mil – Acropora millepora. M. aeq – Montipora aequituburculata*.

|  | May |  |  | June |  |  |
| --- | --- | --- | --- | --- | --- | --- |
|  | No. cultures | Fert. range (%) | Total larvae | No. cultures | Fert. range (%) | Total larvae |
| **2022** |  |  |  |  |  |  |
| *A. aus* | - | - | - | - | - | - |
| *A. els* | 1 <sup>[3]</sup> | 32 <sup>[3]</sup> | 1.0 x10<sup>4</sup> | 1 <sup>[3]</sup> | ND | 1.3 x10<sup>4</sup> |
| *A. hya* | - | - | - | 1 <sup>[3]</sup> | ND | 1.5 x10<sup>4</sup> |
| *A. lor* | - | - | - | 5 <sup>[3-13]</sup> | 65 <sup>[3]</sup> – 77 <sup>[9]</sup> | 8.2 x10<sup>4</sup> |
| *A. mil* | 1 <sup>[2]</sup> | 80 <sup>[2]</sup> | 1.5 x10<sup>4</sup> | 1 <sup>[2]</sup> | ND | 1.5 x10<sup>4</sup> |
| *M. aeq* | - | - | - | - | - | - |
| **2023** |  |  |  |  |  |  |
| *A. aus* | 1 <sup>[3]</sup> | ND | 2.5 x10<sup>4</sup> | - | - | - |
| *A. els* | 1 <sup>[2]</sup> | ND | 5.0 x10<sup>4</sup> | 1 <sup>[3]</sup> | 76 <sup>[3]</sup> | 6.6 x10<sup>4</sup> |
| *A. hya* | - | - | - | 1 <sup>[3]</sup> | 83 <sup>[3]</sup> | 3.1 x10<sup>4</sup> |
| *A. lor* | - | - | - | 3 <sup>[9-17]</sup> | 84 <sup>[9]</sup> - 92 <sup>[17]</sup> | 1.5 x10<sup>5</sup> |
| *A. mil* | 1 <sup>[2]</sup> | ND | 2.5 x10<sup>4</sup> | 2 <sup>[2]</sup> | 78 <sup>[2]</sup> | 2.3 x10<sup>4</sup> |
| *M. aeq* | 2 <sup>[3-6]</sup> | 11 <sup>[3]</sup> - 91 <sup>[6]</sup> | 1.2 x10<sup>6</sup> | 2 <sup>[3-5]</sup> | 94 <sup>[3]</sup> | 2.1 x10<sup>5</sup> |

\*Superscript brackets denote number of individual genotypes pertaining to sample. Total larvae produced represents the sum of all larvae produced from cultures, including those where fertilisation percentage was not determined (ND)

Between 10^th^ and 17^th^ June 2022 (4-11 NAFM) five species spawned, including 28 colonies of *A. loripes*, four *A. hyacinthus*, two *A. millepora*, three *A.elseyi*, and a single colony of *M. aequituberculata*. Like the spawning event in May, some coral colonies spawned partially over multiple nights (Table S1). A total of five larval cultures were established for *A. loripes* over five nights (4-9 NAFM) with contributions of gametes from between 3-13 colonies while a single larval culture was created for each of: *A. elseyi*, *A. millepora* and *A. hyacinthus* using gametes from 2-3 colonies (Table 1). In total, approximately 125,000 larvae were produced in June 2022 (Table 1).

Notably, in 2022 all coral species spawned within the expected NAFM range from GBR observations as reported in Baird et al., (2021) (Fig. 5a), except for *M. aequitub*er*culata* which spawned slightly later (6-10 NAFM) than previous observations (2-6 NAFM) (Fig. 5a). During spawning nights, the corals generally spawned at times closely matching those reported in Baird et al., (2021), especially for those species represented by a large number of observations in the database, such as *A. millepora* (Fig. 5b). A notable exception was that of *M. aequituberculata* which spawned nearly an hour later than previously reported (Fig. 5b), although it should be noted that only two observations exist for this species within the spawning database (Baird et al., 2021).

**Figure 5:**
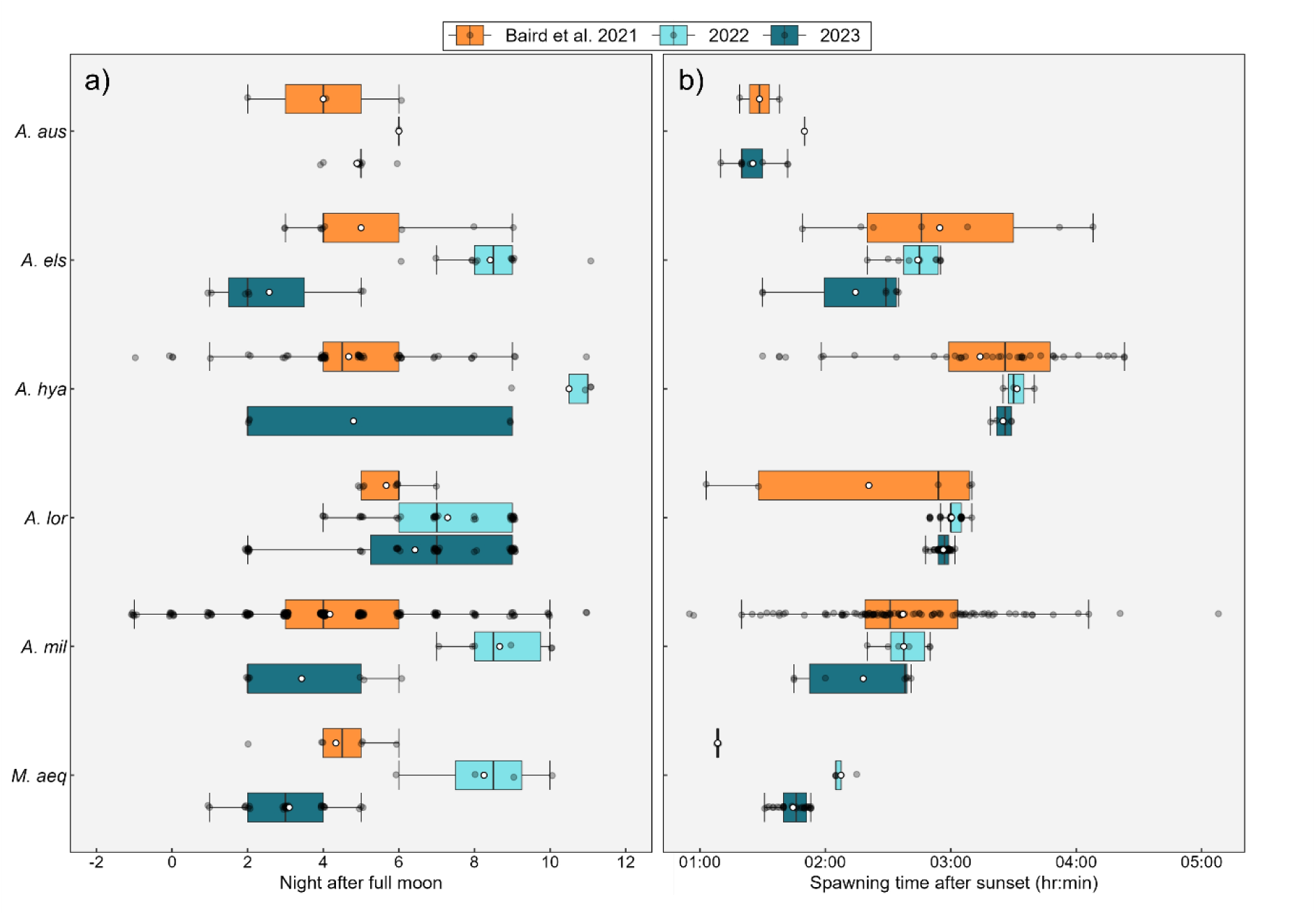
Spawning synchronicity of corals in the out-of-season spawning program compared to wider Great Barrier Reef (GBR) observations. Data shown represent a) spawning night after full moon (NAFM), and b) time after sunset (TAS) data for each species in 2022 and 2023, together with observations from the Indo-Pacific coral spawning database (Baird et al., 2021) corresponding to the Great Barrier Reef (GBR) ecoregion and from the corresponding geomorphological reef group (i.e., inshore or offshore) matching the broodstock (F0) site of origin. NAFM observations comprise: *Acropora elseyi* (n = 9), *Acropora hyacinthus* (n = 28), *Acropora loripes* (n = 9), *Acropora millepora* (n = 43), *Montipora aequituberculata* (n = 6) and *Acropora austera* (n = 3, where both – mid-shelf and inshore reef observations were included due to the small sample size). N.b. n = 3 observations of NAFM (-14 NAFM) were excluded for *A. elseyi* as we were unable to reconcile values reported in the Baird et al., (2021) database with the original source. TAS observations comprise: *A. austera* (n = 2), *A. elseyi* (n = 7), *A. hyacinthus* (n = 20), *A.loripes* (n = 5), *A. millepora* (n = 21) and *M. aequituberculata* (n = 2). Species key: *A. aus* – *Acropora austera. A. els* – *Acropora elseyi. A. hya* – *Acropora hyacinthus. A. lor* – *Acropora loripes. A. mil* – *Acropora millepora. M. aeq* – *Montipora aequituburculata*.

The proportion of F1 broodstock colonies spawning was highest for *A. loripes* and *A. millepora* (75%), followed by *A. hyacinthus* and *A. elseyi* (57.1%), *M. aequituberculata* (16.7%) and *A. austera* (2.6%) (Fig. 2). It was possible to establish larval cultures for four of the six coral species that spawned (*A. millepora, A. elseyi*, *A. loripes,* and *A. hyacinthus*), allowing us to produce a total of approximately 150,000 larvae (Table 1). These larvae were subsequently settled, therefore allowing us to close the life cycle of these four *Acropora* species in captivity within SeaSim (Fig. 4 h-j).

#### 2023

A greater number of individual colonies (59 of 108) spawned in 2023, compared to the previous year (Fig. 2). As in 2022, spawning was split across May and June, with the majority (50 of 59) of colonies spawning in June. Between 3^rd^ May – 7^th^ May 2023 (2-6 NAFM) a total of 16 coral colonies across four species spawned (Table S1). Most corals exhibited partial spawning over multiple nights (Table S1). Fertilisation was successful for all species that spawned, with fertilisation success the highest for *M. aequituberculata* at 91% (Table 1, Fig. 4g). A total of 1.27 million larvae were produced in the May spawning event, the majority of which were from *M. aequituberculata* (1.17 million larvae).

Between 31^st^ May – 8^th^ June 2023 (1-9 NAFM) fifty colonies from six species were observed to spawn (Table S1); with some individuals spawning partially across multiple nights. Successful fertilisation was observed for all species, with *M. aequituberculata* exhibiting the highest fertilisation success of 94% (Table 1). Fertilisation for all *Acropora* species ranged between 76% (*A. elseyi)* and 92% (*A. loripes*) (Table 1). Approximately 475,000 coral larvae were produced in the June spawning event (Table 1).

Across the May and June spawning events in 2023, *A. austera* and *M. aequituberculata* (Fig. 4k) larvae were successfully settled, thus closing the life cycle of these species in captivity. As in 2022, most corals spawned within the expected NAFM and spawning time (hours after sunset) range based on previous GBR observations (Baird et al., 2021) (Fig. 5). Similar to 2022, *M. aequituberculata* spawned approximately 30 minutes later than previously recorded (Baird et al., 2021). The proportion of colonies spawning from the F1 broodstock was highest in *M. aequitub*e*rculata* (100%) followed by *A. loripes* (83%), *A. millepora* (75%), *A. hyacinthus* and *A. elseyi* (both 71%) and *A. austera* (18%) (Fig. 2). Across the six species that spawned in 2023, approximately 1.6 million coral larvae were produced representing an 11-fold increase compared to 2022 (Table 1).

In assessing how closely synchronised coral spawning was between years (2022 - 2023), ΔNAFM values were statistically indistinguishable for most coral species (GLM, p > 0.05, Fig. 5a), except *M. aequitub*e*rculata* (GLM, p < 0.001, Fig. 5a) which exhibited a smaller ΔNAFM in 2023 (i.e., spawning closer to full moon) compared to 2022 (ΔNAFMs 1.1 and 4.3 respectively). Both *A. millepora* and *M. aequituburculata* spawned slightly earlier post-sunset in 2023 compared to 2022 (GLM, p < 0.05, Fig. 5b), while other coral species exhibited no difference in time after sunset (ΔTAS) values between years (GLM, p > 0.05, Fig. 5b).

## Discussion

Research on early coral life stages often requires access to coral gametes, larvae and recruits. However, since many coral species are annual broadcast spawners, the availability of such biological material is restricted to a narrow time window during the year and imposes a problematic bottleneck to research programs. In this study, we utilise advanced aquarium technology to replicate and offset seasonality of environmental cues (temperature, photoperiod and moonlight), inducing predictable out-of-season spawning of GBR corals in austral winter - successfully generating >2 million coral larvae. We discuss the physiological insights gained surrounding environmental regulation of coral spawning and how breeding corals out of season could facilitate and accelerate research efforts into early life history and processes of corals.

### Insights into environmental regulation of spawning synchronisation

Corals maintained under seasonally-offset environmental conditions spawned six months apart from their natural reef counterparts as expected (i.e., May/June rather than November/December). The timing of spawning closely matched literature observations of natural mass spawning events, both in terms of nights after full moon (NAFM) and time-of-day (e.g., Babcock et al., 1986; Baird et al., 2021). These findings add to the growing body of evidence (Brady et al., 2009; Keith et al., 2016; Craggs et al., 2020; Sakai et al., 2024) suggesting temperature, sunlight and moonlight are the primary environmental cues regulating the synchronisation of coral mass spawning events. Although we did not simulate tidal patterns within our aquaria systems, spawning was still observed close to previous literature reports, suggesting that while tidal patterns may be correlative with those cues for spawning synchronicity, they may have minimal direct effect on the coral taxa examined here. However, we are unable to exclude the possibility that location-specific conditions - such as site of origin - may have contributed to this observed outcome. Further development of the system described here could of course allow for greater manipulation of these parameters to better understand endogenous (circadian rhythms) and exogenous controls that drive spawning synchrony in corals.

Interestingly, pigmented eggs were observed in two of the larger coral colonies in June 2020, less than 12 months after the offset temperature profile was initiated. This supports observations that temperature change is strongly linked to onset of gametogenesis and suggests capacity for some corals to synchronise to a modified temperature profile inside a year. During the following year (2021), offset day length and moonlight profiles were incorporated into the system. Subsequently, a total of eight coral colonies from three species were observed to have pigmented eggs one week prior to expected spawning dates, again suggesting corals were synchronised to the modified (offset) environmental profile. However, no gamete release was observed for any taxa over the expected spawning period (i.e., 0 - 11 NAFM, June/July) – presumably due to aforementioned issues with moonlight profile programming (see Materials and Methods). While this outcome was unfortunate for the experiment, it did appear to reinforce the importance of moonlight within the hierarchy of cues coordinating synchronous gamete release (e.g., Kaniewska et al., 2015; Komoto et al., 2023; Sakai et al., 2024). Nightly irradiance from moonlight modulates circadian rhythms in corals (Jokiel et al., 1985; Brady et al., 2016) and lunar periodicity has long been recognised in studies of coral reproduction. Moonlight may also be regulating synchronicity in corals at finer temporal scales. It is plausible, as found in some other marine invertebrates, e.g., Zurl et al., (2022), that diurnal exposure to moonlight may aid in regulating the precise timing of spawning in corals. It is also important to note that we were unable to disentangle the contribution of time (i.e., coral age) when comparing spawning observations between years, so this should also be considered in the interpretation of the results presented in this study.

In contrast to the evidence suggesting moonlight was important for regulation of spawning synchronisation in this study, the importance of solar insolation appears more equivocal. Although sunrise and sunset times were dynamically profiled to match seasonal changes in photoperiod, maximum daily light intensities were kept constant throughout the experiment, with a peak intensity of 150 µmol photons m^-2^ s^-1^ (matching their long-term growth history). As such, the cumulative daily photon dose (or daily light integral) experienced by corals in this study was lower, and exhibited less seasonal variance than corals would typically experience on a natural reef system (see Anthony et al., 2004; Strahl et al., 2019; Noonan et al., 2022). Therefore, while it appears that full replication of seasonal insolation variance is not needed to achieve predictable spawning in the coral taxa here - and thus may be relatively unimportant as an environmental cue – we acknowledge that it is not possible to tease apart any effect(s) of photoperiod modulation. Certainly, insolation is thought to serve as an important spawning cue in low-latitude reefs where annual SST variability is inherently lower than tropical reef systems (Penland et al., 2004) and where its importance as a cue in higher-latitude reef systems such as the GBR is presumably overshadowed by that of SST variance.

Out-of-season spawning may also provide a useful platform to examine environmental factors contributing to asynchronisation of coral spawning (Fig. 6). For example, artificial light at night (ALAN) is an emerging threat to coastal reef systems, altering diel cycles of organisms including corals, modifying coral polyp feeding behaviour (Mardones et al., 2023) and causing delayed gametogenesis among key broadcast spawning coral species (Kaniewska et al., 2015; Neely et al., 2020; Ayalon et al., 2021; Davies et al., 2023). To date, knowledge of how ALAN may disrupt spawning cues in reef organisms is largely based on observational studies, yet out-of-season spawning programs provide a clear manipulative way to test such effects on corals *ex-situ*.

**Figure 6:**
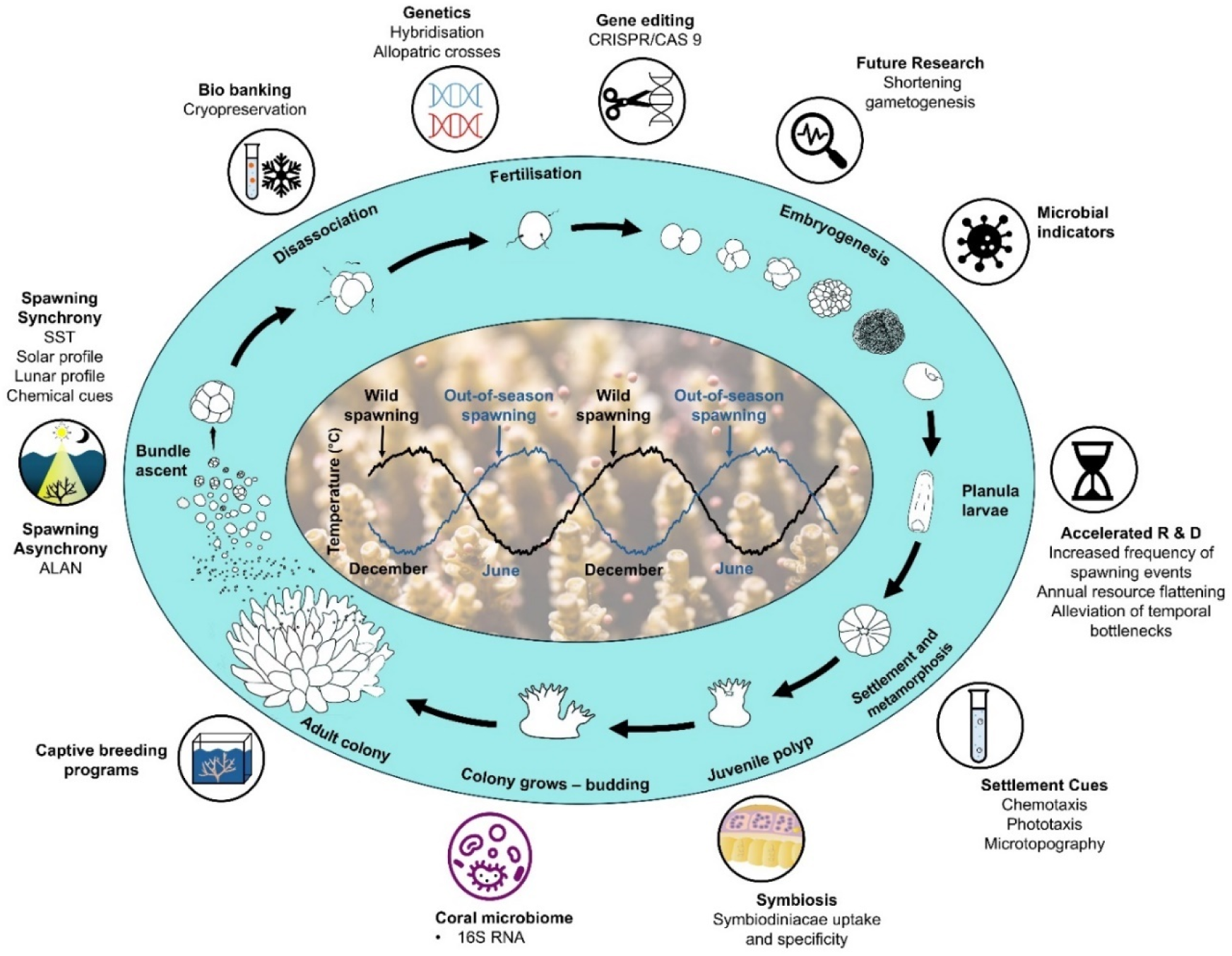
Conceptual figure outlining potential applications for out-of-season coral spawning programs including a) acceleration of existing research areas (e.g., gene editing, biobanking, coral-algal symbiosis establishment, larval settlement cues and inhibitors), b) novel applications (e.g., coral hybridisation opportunities, shortening coral gametogenesis cycle), c) as an experimental platform to test spawning cues (chemical cues, lunar profile, solar profile, sea surface temperature [SST], artificial light at night [ALAN]) at the various/relevant coral life stages (indicated by the shaded blue area - adapted from Jones et al., 2015; Randall et al., 2020).

### A tool to accelerate research into coral early life history stages

Out-of-season spawning can accelerate research efforts (Fig. 6) by theoretically allowing year-round access to study coral early life stages and processes, limited primarily by logistics (space, cost and, effort), rather than fundamental organismal biology *per se*. However, the nature and extent to which coral larvae produced *ex-situ* out-of-season can substitute larvae from natural mass spawning events for diverse research applications, particularly ecophysiological studies (e.g., Alderdice et al., 2022), will depend on understanding any physiological difference(s) that may exist between them. Previous work has shown that various aspects of larval physiology (developmental timing, energy reserves, fertilisation success, competency, recruitment success and growth) are influenced by environmental history and impacted by environmental stress (e.g., Edmunds et al., 2001; Negri et al., 2007; Rivest et al., 2017). Whether synchronisation to offset environmental profiles confers additional physiological stress to the coral that subsequently impacts larval quality is unknown and represents a critical area for further investigation.

While understanding how *ex-situ* offset environmental profiles impact coral larval fitness is clearly an important future step, there are several applications where use of “out-of-season” coral larvae could provide an immediate research boost (Fig. 6). For example, CRISPR/Cas9 genome editing - which has been applied to coral zygotes via microinjection techniques - is highly time- and labour-intensive (Cleves et al., 2020), therefore a longer time window for zygote availability would likely catalyse significant advances in such genetics and gene-function studies. The possibility of year-round access to coral gametes could also facilitate advancements in other research areas, for example, supporting workflows and processes aimed at understanding (and developing) biological and chemical cues for inducing coral larval settlement (e.g., Abdul Wahab et al., 2023). Shortening the gametogenesis cycle through further manipulation of environmental profile(s) could also reduce the time window between captive spawning and represents an interesting but, as yet unexplored, research avenue. It would, however, be prudent to explore any associated physiological trade-offs (e.g., Moreno, 1993) in parallel.

### Towards an improved understanding of coral husbandry and nutrition

Here we report long-term success in maintaining key GBR corals in a controlled aquarium environment, including the closure of their life cycle. While species-specificity likely exists in GBR corals captive requirements (Borneman, 2000), the aquarium conditions and nutritional regimes applied in this study provide a strong foundation for the long-term holding of diverse GBR coral taxa.

There inevitably exists further scope for improvement however, as evidenced by occasional neutral buoyancy in egg/sperm bundles released by *A. loripes* in this study; a phenomenon rarely reported in nature for this species. Positive buoyancy is a key trait to ensure optimal fertilisation and dispersal of egg/sperm bundles (Harii et al., 2002, 2007; Ricardo et al., 2016; Okubo et al., 2020), and wax esters - a type of lipid involved in energy storage - are the main contributor to positive buoyancy in coral eggs (Arai et al., 1993; Okubo et al., 2020). Previous examinations of tissue macromolecular composition in corals both before and after spawning further show that most of the energy invested into coral reproduction originates from lipid metabolism (Leuzinger et al., 2003). Although lipid content and composition of gametes was not explicitly examined here, previous work suggests heterotrophic feeding (particle capture) and high light exposure drives increased lipid content in corals (Al-Moghrabi et al., 1995; Oku et al., 2003; Zhukova and Titlyanov, 2006; Treignier et al., 2008). Here, we provided corals with a selection of heterotrophic feeds (microalgae, rotifers, and *Artemia salina*), ranging from ∼15 –750 µm in size to accommodate the variable polyp sizes found across the six species in this experiment (∼ 0.4 - 1.6 mm, Madin et al., 2016). However, understanding what constitutes an “optimal” feeding regime for captive corals is continually evolving (e.g., Conlan et al., 2018, 2019; Saper et al., 2023) and we are unable to exclude the possibility that decreased buoyancy of egg/sperm bundles observed in *A. loripes* was linked to nutritional deficiencies.

In addition to heterotrophy, light has been identified as a key limiting resource governing energy investment into reproduction for reef-forming corals (Leuzinger et al., 2012). Specifically, Leuzinger et al., (2012) found that partial shading (equivalent to ∼35 % incident irradiance) of *Montipora digitata* colonies at Heron Island (Southern GBR) caused decreased energy investment into reproduction compared to unshaded corals, while colonies subjected to full-shading (∼3 % incident irradiance) halted gametogenesis altogether. Thus, energy allocation strategies in corals are inherently flexible and depend on the amount of light relative to the corals’ photoacclimation status (Hennige et al., 2008). In this study, light levels provided were clearly sufficient to facilitate allocation of energy towards reproduction for all six coral species examined, but whether potential differences in energy allocation strategies resulting from differences in photoacclimation status may have contributed to observed neutral buoyancy in *A. loripes* egg/sperm bundles is unknown. Likely, the total daily photon dose provided (∼5.5 mol photons m^-2^ d^-1^) was lower than that available to corals *in-situ* at their respective depth and site of origin (e.g., see Noonan et al., 2022), so it is plausible that energetic resources invested into reproduction for *A. loripes* (and indeed for the other coral species examined) could be boosted by increased irradiance.

### Out-of-season spawning – viability as a tool to support reef restoration efforts

Global degradation of coral ecosystems has resulted in a surge in active reef restoration initiatives aimed at supplementing traditional passive habitat protection measures (Boström-Einarsson et al., 2020). Outplanting of corals onto degraded reefs is a strategy routinely employed to increase coral cover and enhance reef architectural complexity (Boström-Einarsson et al., 2020). While many outplanting programmes rely on asexual propagation, or “fragments of opportunity”, from local coral colonies *in-situ*, use of sexually propagated coral recruits into reef restoration programs can enhance genetic diversity to boost climate change resilience (Barton et al., 2017). Therefore, coral aquaculture - the method of raising large numbers of corals in captive aquaria - is increasingly viewed as a key component of the reef restoration toolbox (Leal et al., 2016; Barton et al., 2017).

To date, a role for coral aquaculture has involved collection of gametes which are fertilised, settled onto artificial structures and reared *ex-situ* before outplanting (e.g., Omori, 2005), or using gametes that are collected and fertilised in temporary holdings and released in a controlled fashion onto predetermined reef areas (e.g., Cruz and Harrison, 2017). Out-of-season spawning necessitates holding of broodstock year-round in aquaria with sophisticated environmental control which may impose techno-economic constraints for using such larvae in outplanting efforts. Specifically, the initial cost of aquarium equipment and ongoing maintenance, energy costs to maintain temperature against ambient (external) conditions, and provision of year-round coral husbandry impacts cost-effectiveness. On the other hand, out-of-season spawning programs allow for outplanting of coral recruits at different times of the year (pending successful acclimation to ambient temperature), where physicochemical (e.g., light, currents, sediments) and biological (e.g., planktonic supply, corallivory) selective pressures might be more favourable and hence boost relative return on effort (RRE, Suggett et al., 2019) through increased survivorship. Growing evidence suggests that epigenetic changes arising from exposure to favourable biological and physical conditions at specific coral nurseries during early-life stages enhance corals’ ability to endure environmental perturbation (Rinkevich, 2019) – but whether similar benefits could be derived by outplanting coral recruits out of season remains an open question. With coral cryopreservation technologies advancing at a rapid pace (Daly et al., 2018), out-of-season spawning programs provide a platform for testing the viability of outplanting (reanimated) larvae outside of natural spawning windows as part of future possible restoration efforts (Hagedorn et al., 2019).

### Future directions

This study marks the first successful demonstration of “out-of-season” spawning for GBR corals through controlled manipulation of environmental cues in aquaria. Ability to generate coral gametes and larvae “on demand” has the potential to accelerate existing research programs, particularly regarding reproductive ecology and active reef interventions, while also creating opportunities for novel research (Fig. 6). For example, the ability to precisely control the time of spawning for specific coral species raises the intriguing possibility of exploring hybridisation opportunities between corals that would otherwise remain temporally separated by differences in spawning time(s) and date(s). This could provide an opportunity to create novel coral phenotypes (e.g., Suggett et al., 2019), potentially with outbreeding enhancement (or hybrid vigour, see Goulet et al., 2017; Lamb et al., 2024) that could boost local reef resilience if outplanted onto degraded reef areas.

Out-of-season spawning represents an underexplored technology for corals, with a number of novel applications likely to emerge over time. As such, we expect out-of-season spawning to form a key part of the growing toolbox of active restoration strategies aimed at reversing the global decline of coral reefs.

## Acknowledgements

We acknowledge valuable discussions across the peer community in refining the methodology and concepts presented here; in particular, we thank Dr Jamie Craggs (Project Coral). This work was facilitated by the considerable commitment and efforts of multiple SeaSim staff. Here, we would like to thank Steven Green for assistance with controls, Shanae Read, Andrea Dobrescu, Dr Jonathan Barton, Natalie Giofre, Stephanie Garra, and Sara Godinez for various aspects of coral husbandry. Jonathan Armitage, Anton Rocconi, Samantha Jaworski and Christine Giuliano also provided valuable coral husbandry during the crucial early life stage (F1), while Justin Hochen, Grant Milton, Eduardo Arias, Tom Barker, and Tristan Lever provided technical support with experimental setup, maintenance, and troubleshooting. Thanks also to Dr Gerard Ricardo for providing insightful comments which substantially improved the first draft of this manuscript.

## Notes

### Competing Interest Statement

The authors have declared no competing interest.

