## Supplementary Figures for "Controlled “out-of-season” spawning of reef-forming corals in aquaria using offset environmental profiles"

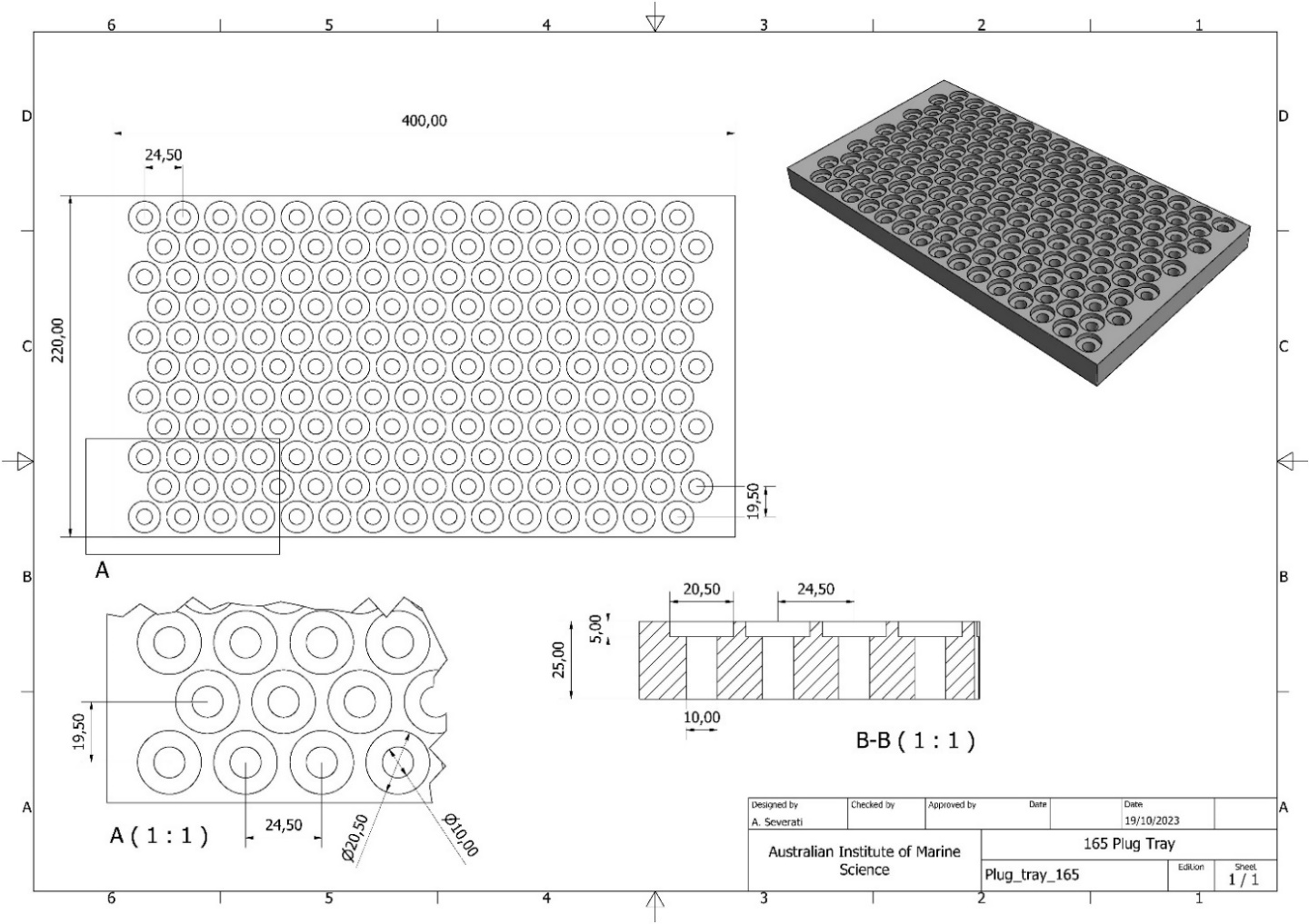

**Figure S1.** Custom made Poly Vinyl Chloride trays used to hold 165 x Ocean Wonders 20mm diameter aragonite frag plugs for coral larvae settlement in the out-of-season spawning project.

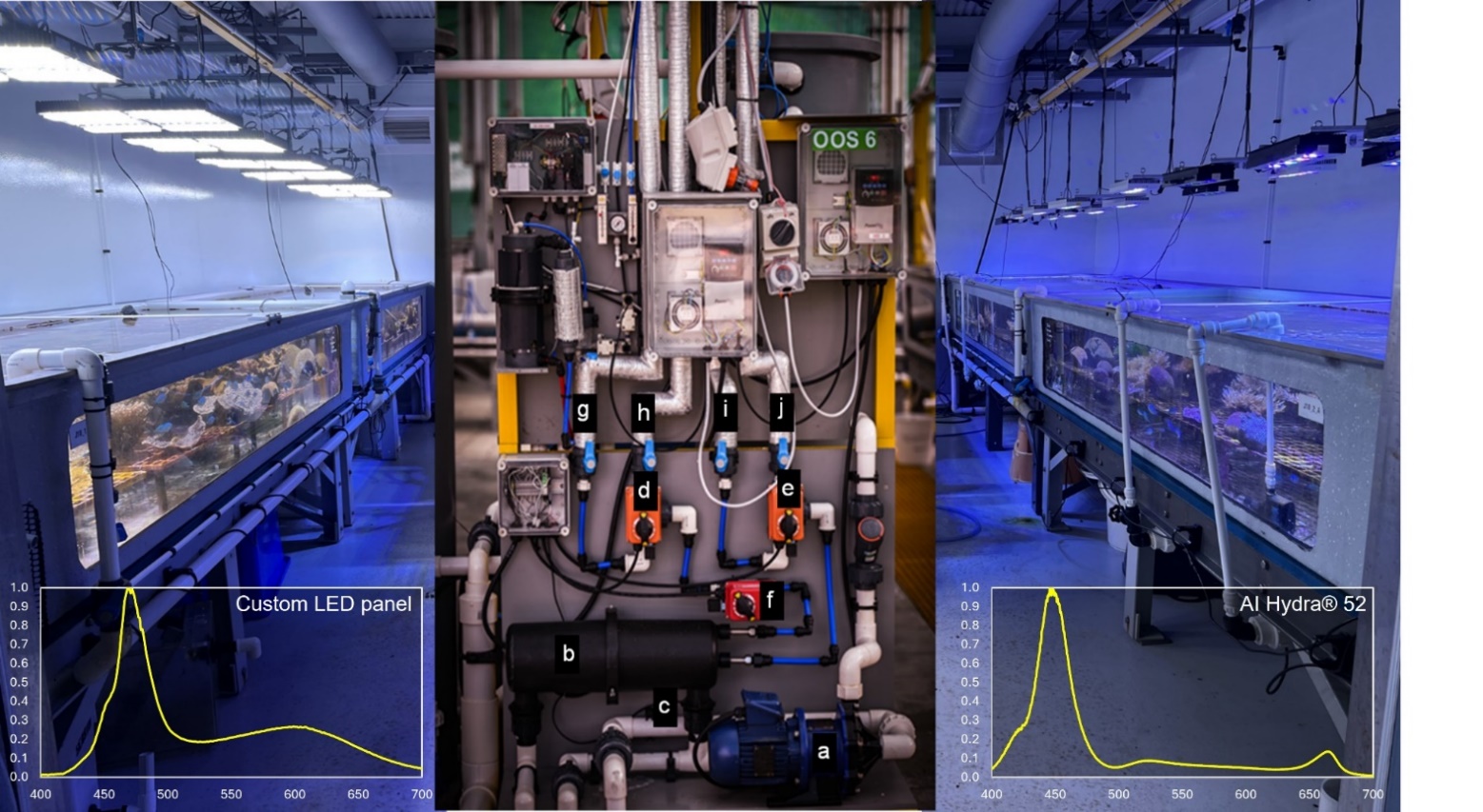

**Figure S2.** Lighting and life support systems for the F1 coral holding system. a) Metallic centrifugal magnetic drive pump b) Shell and tube heat exchanger c) FEP insulated resistance temperature detector d) Electric actuating valve controlling cold (15°C) or hot (40°C) water delivery and e) return f) Electric actuating valve controlling percentage ( 0-100 %) of water delivery through HEX for system temperature control g) Cold (15°C) water delivery h) Hot (40°C) water delivery i) Cold (15°C) water return j) Hot (40°C) water return. Inset panels show spectral output of the two lighting types used (See Materials and Methods).

NO3- (µmol/L)

|  | J18A | J18B |
| --- | --- | --- |
| 10/06/2020 | 0.805 | 0.485 |
| 22/06/2020 | 1.475 | 0.38 |
| 1/07/2020 | 0.925 | 0.705 |
| 8/07/2020 | 1.015 | 0.595 |
| 15/07/2020 | 0.99 | 0.535 |
| 22/07/2020 | 0.805 | 0.315 |
| 29/07/2020 | 0.085 | 0.15 |
| 5/08/2020 | 0.13 | 0.165 |
| 12/08/2020 | 0.04 | 0.13 |
| 20/08/2020 | 0.485 | 0.15 |
| 26/08/2020 | 0.58 | 0.245 |
| 2/09/2020 | 0.42 | 0.045 |
| 9/09/2020 | 0.465 | 0.14 |
| 16/09/2020 | 0.175 | 0.045 |
| 23/09/2020 | 0.415 | 0.125 |
| 2/10/2020 | 0.505 | 0.07 |
| 7/10/2020 | 0.485 | 0.105 |
| 14/10/2020 | 0.585 | 0.08 |
| 21/10/2020 | 0.685 | 0.09 |
| 4/11/2020 | 1.045 | 0.185 |
| 12/11/2020 | 1.185 | 0.21 |
| 19/11/2020 | 1.27 | 0.605 |
| 24/11/2020 | 1.205 | 0.36 |
| 3/12/2020 | 1.28 | 0.77 |
| 9/12/2020 | 1.64 | 1.305 |
| 16/12/2020 | 1.875 | 0.735 |
| 23/12/2020 | 1.22 | 0.72 |
| 30/12/2020 | 1.36 | 1.205 |
| 6/01/2021 | 1.835 | 1.05 |
| 13/01/2021 | 2.425 | 1.37 |
| 20/01/2021 | 2.55 | 1.515 |
| 27/01/2021 | 2.66 | 1.185 |
| 3/02/2021 | 1.91 | 1.33 |
| 10/02/2021 | 1.665 | 1.04 |
| 17/02/2021 | 1.99 | 1.405 |
| 24/02/2021 | 2.525 | 1.305 |
| 3/03/2021 | 2.38 | 1.65 |
| 10/03/2021 | 1.56 | 0.82 |
| 16/03/2021 | 2.525 | 1.425 |
| 24/03/2021 | 2.545 | 1.28 |
| 31/03/2021 | 2.625 | 1.325 |
| 7/04/2021 | 1.9 | 1.255 |
| 14/04/2021 | 1.735 | 1.7 |
| 21/04/2021 | 1.97 | 1.84 |
| 28/04/2021 | 2.37 | 3.22 |
| 5/05/2021 | 1.625 | 1.84 |
| 12/05/2021 | 1.845 | 1.355 |
| 19/05/2021 | 1.785 | 1.18 |
| 26/05/2021 | 1.135 | 0.965 |
| 2/06/2021 | 1.65 | 1.055 |
| 9/06/2021 | 2.045 | 1.195 |
| 16/06/2021 | 1.11 | 0.745 |
| 23/06/2021 | 1.045 | 2.105 |
| 30/06/2021 | 1.18 | 0.83 |
| 7/07/2021 | 1.215 | 0.825 |
| 14/07/2021 | 1.25879 | 0.923654 |
| 21/07/2021 | 1.415 | 0.795 |
| 27/07/2021 | 1.34 | 0.71 |
| 4/08/2021 | 1.86 | 0.965 |
| 11/08/2021 | 0.675 | 1.265 |
| 18/08/2021 | 1.18 | 1.375 |
| 25/08/2021 | 1.36 | 1.51 |
| 1/09/2021 | 1.4 | 1.59 |
| 8/09/2021 | 1.15 | 1.38 |
| 15/09/2021 | 1.085 | 1.39 |
| 21/09/2021 | 1.15 | 1.45 |
| 29/09/2021 | 1.1 | 2.165 |
| 6/10/2021 | 1.135 | 1.625 |
| 13/10/2021 | 0.93 | 1.4 |
| 20/10/2021 | 0.88 | 0.915 |
| 27/10/2021 | 1.05 | 0.71 |
| 3/11/2021 | 0.985 | 0.935 |
| 10/11/2021 | 1.22 | 1.24 |
| 17/11/2021 | 1.615 | 1.095 |
| 24/11/2021 | 1.285 | 1.34 |
| 1/12/2021 | 1.145 | 1.4 |
| 8/12/2021 | 0.97 | 1.11 |
| 15/12/2021 | 1.145 | 1.36 |
| 22/12/2021 | 1.415 | 1.21 |
| 28/12/2021 | 0.935 | 1.215 |
| 5/01/2022 | 1.36 | 1.2 |
| 12/01/2022 | 1.755 | 1.685 |
| 19/01/2022 | 1.895 | 1.73 |
| 25/01/2022 | 1.755 | 1.86 |
| 2/02/2022 | 2.015 | 2.125 |
| 9/02/2022 | 1.945 | 1.91 |
| 16/02/2022 | 2.415 | 2.665 |
| 23/02/2022 | 2.37 | 2.605 |
| 2/03/2022 | 2.495 | 3.24 |
| 9/03/2022 | 1.99 | 1.495 |
| 16/03/2022 | 1.845 | 1.71 |
| 23/03/2022 | 1.925 | 1.87 |
| 30/03/2022 | 1.925 | 2.045 |
| 6/04/2022 | 2.445 | 2.32 |
| 13/04/2022 | 2.845 | 2.67 |
| 19/04/2022 | 2.475 | 2.575 |
| 27/04/2022 | 3.955 | 4.68 |
| 4/05/2022 | 5.36 | 9.41 |
| 11/05/2022 | 4.03 | 5.735 |
| 19/05/2022 | 4.355 | 4.595 |
| 25/05/2022 | 4.69 | 4.18 |
| 1/06/2022 | 3.8 | 3.165 |
| 8/06/2022 | 3.805 | 3.225 |
| 15/06/2022 | 3.32 | 3.055 |
| 24/06/2022 | 2.82 | 2.23 |
| 29/06/2022 | 2.515 | 2.69 |
| 6/07/2022 | 2.425 | 1.51 |
| 13/07/2022 | 2.245 | 1.335 |
| 20/07/2022 | 2.065 | 1.055 |
| 27/07/2022 | 1.985 | 0.625 |
| 3/08/2022 | 1.845 | 1.19 |
| 10/08/2022 | 1.71 | 1.415 |
| 17/08/2022 | 2.345 | 1.485 |
| 24/08/2022 | 2.165 | 1.08 |
| 31/08/2022 | 2.135 | 1.235 |
| 7/09/2022 | 2.33 | 1.145 |
| 14/09/2022 | 2.38 | 1.495 |
| 20/09/2022 | 3.055 | 1.49 |
| 27/09/2022 | 3.075 | 1.445 |
| 5/10/2022 | 2.62 | 0.91 |
| 12/10/2022 | 2.56 | 1.335 |
| 19/10/2022 | 2.455 | 1.515 |
| 26/10/2022 | 2.29 | 1.78 |
| 1/11/2022 | 1.705 | 1.46 |
| 8/11/2022 | 1.52 | 1.12 |
| 15/11/2022 | 2.34 | 1.75 |
| 22/11/2022 | 2.2 | 1.85 |
| 29/11/2022 | 2.425 | 2.16 |
| 6/12/2022 | 2.11 | 1.93 |
| 13/12/2022 | 2.01 | 1.92 |
| 20/12/2022 | 2.495 | 2.205 |
| 27/12/2022 | 1.905 | 1.65 |
| 3/01/2023 | 1.695 | 1.28 |
| 10/01/2023 | 2.645 | 1.295 |
| 17/01/2023 | 3.93 | 1.595 |
| 24/01/2023 | 2.38 | 1.435 |
| 1/02/2023 | 2.125 | 1.53 |
| 8/02/2023 | 2.23 | 1.28 |
| 15/02/2023 | 2.43 | 1.43 |
| 22/02/2023 | 3.28 | 1.7 |
| 1/03/2023 | 4.665 | 2.04 |
| 8/03/2023 | 3.815 | 2.29 |
| 15/03/2023 | 3.17 | 2.115 |
| 22/03/2023 | 2.93 | 2.36 |
| 29/03/2023 | 2.995 | 2.415 |
| 6/04/2023 | 2.935 | 2.49 |
| 11/04/2023 | 2.605 | 2.17 |
| 19/04/2023 | 3.075 | 2.92 |
| 26/04/2023 | 3.26 | 4.055 |
| 3/05/2023 | 4.125 | 4.265 |
| 10/05/2023 | 2.895 | 3.01 |
| 17/05/2023 | 3.1 | 2.66 |
| 24/05/2023 | 2.77 | 2.525 |
| 30/05/2023 | 3.37 | 2.78 |
| 6/06/2023 | 2.895 | 3.03 |
| 14/06/2023 | 3.805 | 4.435 |
| 20/06/2023 | 3.53 | 4.6 |

PO43- (µmol/L)

|  | J18A | J18B |
| --- | --- | --- |
| 10/06/2020 | 0.305 | 0.115 |
| 22/06/2020 | 0.315 | 0.085 |
| 1/07/2020 | 0.205 | 0.2 |
| 8/07/2020 | 0.22 | 0.17 |
| 15/07/2020 | 0.15 | 0.115 |
| 22/07/2020 | 0.14 | 0.085 |
| 29/07/2020 | 0.085 | 0.095 |
| 5/08/2020 | 0.07 | 0.085 |
| 12/08/2020 | 0.06 | 0.09 |
| 20/08/2020 | 0.135 | 0.09 |
| 26/08/2020 | 0.135 | 0.095 |
| 2/09/2020 | 0.15 | 0.05 |
| 9/09/2020 | 0.15 | 0.085 |
| 16/09/2020 | 0.09 | 0.05 |
| 23/09/2020 | 0.125 | 0.045 |
| 2/10/2020 | 0.125 | 0.02 |
| 7/10/2020 | 0.14 | 0 |
| 14/10/2020 | 0.165 | 0 |
| 21/10/2020 | 0.145 | 0 |
| 4/11/2020 | 0.215 | 0.02 |
| 12/11/2020 | 0.235 | 0.06 |
| 19/11/2020 | 0.085 | 0.06 |
| 24/11/2020 | 0.25 | 0.05 |
| 3/12/2020 | 0.135 | 0.09 |
| 9/12/2020 | 0.15 | 0.085 |
| 16/12/2020 | 0.145 | 0.08 |
| 23/12/2020 | 0.12 | 0.095 |
| 30/12/2020 | 0.15 | 0.085 |
| 6/01/2021 | 0.18 | 0.13 |
| 13/01/2021 | 0.275 | 0.205 |
| 20/01/2021 | 0.36 | 0.295 |
| 27/01/2021 | 0.325 | 0.295 |
| 3/02/2021 | 0.365 | 0.26 |
| 10/02/2021 | 0.26 | 0.21 |
| 17/02/2021 | 0.28 | 0.23 |
| 24/02/2021 | 0.335 | 0.215 |
| 3/03/2021 | 0.49 | 0.46 |
| 10/03/2021 | 0.335 | 0.23 |
| 16/03/2021 | 0.35 | 0.265 |
| 24/03/2021 | 0.33 | 0.24 |
| 31/03/2021 | 0.405 | 0.28 |
| 7/04/2021 | 0.305 | 0.27 |
| 14/04/2021 | 0.27 | 0.24 |
| 21/04/2021 | 0.34 | 0.265 |
| 28/04/2021 | 0.46 | 0.3 |
| 5/05/2021 | 0.235 | 0.29 |
| 12/05/2021 | 0.28 | 0.21 |
| 19/05/2021 | 0.35 | 0.23 |
| 26/05/2021 | 0.275 | 0.26 |
| 2/06/2021 | 0.43 | 0.26 |
| 9/06/2021 | 0.42 | 0.295 |
| 16/06/2021 | 0.29 | 0.23 |
| 23/06/2021 | 0.22 | 0.21 |
| 30/06/2021 | 0.315 | 0.235 |
| 7/07/2021 | 0.27 | 0.21 |
| 14/07/2021 | 0.293487 | 0.266596 |
| 21/07/2021 | 0.29 | 0.21 |
| 27/07/2021 | 0.33 | 0.25 |
| 4/08/2021 | 0.365 | 0.225 |
| 11/08/2021 | 0.19 | 0.275 |
| 18/08/2021 | 0.27 | 0.23 |
| 25/08/2021 | 0.295 | 0.25 |
| 1/09/2021 | 0.295 | 0.22 |
| 8/09/2021 | 0.27 | 0.22 |
| 15/09/2021 | 0.24 | 0.225 |
| 21/09/2021 | 0.27 | 0.205 |
| 29/09/2021 | 0.245 | 0.29 |
| 6/10/2021 | 0.23 | 0.245 |
| 13/10/2021 | 0.22 | 0.245 |
| 20/10/2021 | 0.21 | 0.195 |
| 27/10/2021 | 0.255 | 0.15 |
| 3/11/2021 | 0.27 | 0.25 |
| 10/11/2021 | 0.22 | 0.27 |
| 17/11/2021 | 0.17 | 0.155 |
| 24/11/2021 | 0.185 | 0.185 |
| 1/12/2021 | 0.15 | 0.155 |
| 8/12/2021 | 0.14 | 0.135 |
| 15/12/2021 | 0.155 | 0.145 |
| 22/12/2021 | 0.19 | 0.16 |
| 28/12/2021 | 0.07 | 0.105 |
| 5/01/2022 | 0.12 | 0.065 |
| 12/01/2022 | 0.245 | 0.195 |
| 19/01/2022 | 0.19 | 0.17 |
| 25/01/2022 | 0.215 | 0.185 |
| 2/02/2022 | 0.225 | 0.19 |
| 9/02/2022 | 0.215 | 0.17 |
| 16/02/2022 | 0.27 | 0.225 |
| 23/02/2022 | 0.26 | 0.245 |
| 2/03/2022 | 0.235 | 0.235 |
| 9/03/2022 | 0.19 | 0.125 |
| 16/03/2022 | 0.245 | 0.225 |
| 23/03/2022 | 0.25 | 0.2 |
| 30/03/2022 | 0.23 | 0.205 |
| 6/04/2022 | 0.26 | 0.22 |
| 13/04/2022 | 0.23 | 0.185 |
| 19/04/2022 | 0.265 | 0.245 |
| 27/04/2022 | 0.315 | 0.205 |
| 4/05/2022 | 0.41 | 0.36 |
| 11/05/2022 | 0.36 | 0.315 |
| 19/05/2022 | 0.335 | 0.32 |
| 25/05/2022 | 0.325 | 0.225 |
| 1/06/2022 | 0.29 | 0.22 |
| 8/06/2022 | 0.305 | 0.235 |
| 15/06/2022 | 0.285 | 0.235 |
| 24/06/2022 | 0.295 | 0.18 |
| 29/06/2022 | 0.285 | 0.28 |
| 6/07/2022 | 0.3 | 0.245 |
| 13/07/2022 | 0.315 | 0.22 |
| 20/07/2022 | 0.26 | 0.21 |
| 27/07/2022 | 0.26 | 0.17 |
| 3/08/2022 | 0.37 | 0.275 |
| 10/08/2022 | 0.23 | 0.245 |
| 17/08/2022 | 0.28 | 0.22 |
| 24/08/2022 | 0.27 | 0.19 |
| 31/08/2022 | 0.28 | 0.205 |
| 7/09/2022 | 0.28 | 0.25 |
| 14/09/2022 | 0.28 | 0.23 |
| 20/09/2022 | 0.31 | 0.28 |
| 27/09/2022 | 0.295 | 0.21 |
| 5/10/2022 | 0.29 | 0.18 |
| 12/10/2022 | 0.265 | 0.2 |
| 19/10/2022 | 0.27 | 0.2 |
| 26/10/2022 | 0.25 | 0.21 |
| 1/11/2022 | 0.245 | 0.2 |
| 8/11/2022 | 0.31 | 0.3 |
| 15/11/2022 | 0.255 | 0.205 |
| 22/11/2022 | 0.215 | 0.19 |
| 29/11/2022 | 0.21 | 0.195 |
| 6/12/2022 | 0.245 | 0.22 |
| 13/12/2022 | 0.235 | 0.225 |
| 20/12/2022 | 0.325 | 0.305 |
| 27/12/2022 | 0.24 | 0.255 |
| 3/01/2023 | 0.22 | 0.225 |
| 10/01/2023 | 0.24 | 0.17 |
| 17/01/2023 | 0.09 | 0.025 |
| 24/01/2023 | 0.445 | 0.365 |
| 1/02/2023 | 0.37 | 0.325 |
| 8/02/2023 | 0.325 | 0.23 |
| 15/02/2023 | 0.29 | 0.205 |
| 22/02/2023 | 0.355 | 0.255 |
| 1/03/2023 | 0.38 | 0.285 |
| 8/03/2023 | 0.32 | 0.27 |
| 15/03/2023 | 0.32 | 0.22 |
| 22/03/2023 | 0.31 | 0.275 |
| 29/03/2023 | 0.31 | 0.28 |
| 6/04/2023 | 0.335 | 0.3 |
| 11/04/2023 | 0.3 | 0.25 |
| 19/04/2023 | 0.31 | 0.285 |
| 26/04/2023 | 0.365 | 0.415 |
| 3/05/2023 | 0.375 | 0.375 |
| 10/05/2023 | 0.29 | 0.29 |
| 17/05/2023 | 0.335 | 0.28 |
| 24/05/2023 | 0.3 | 0.25 |
| 30/05/2023 | 0.315 | 0.27 |
| 6/06/2023 | 0.3 | 0.335 |
| 14/06/2023 | 0.355 | 0.38 |
| 20/06/2023 | 0.34 | 0.35 |

**Figure S3**. Concentrations of dissolved nutrients (nitrate (NO_3_^-^), nitrite, ammonia, and phosphate (PO_4_^3-^), for both broodstock holding systems measured over the duration of the study.
