## Supplementary Methods for "Controlled “out-of-season” spawning of reef-forming corals in aquaria using offset environmental profiles"

**Supplementary methods - Closing the life cycle of captive coral species (F2 generation)**

To close the lifecycle of the coral species in the out-of-season spawning program, larvae were settled during May and June spawning events in both 2022 and 2023 following methodology as described in Materials & Methods. Briefly, second filial generation (F2) coral recruits were held in 50 L aquaria supplied with filtered (1 µm) seawater providing one system turnover per hour. F2 coral recruits were inoculated seven days post-settlement with cultured photosynthetic endosymbionts (Family: Symbiodiniaceae, LaJeunesse et al. 2018) specifically: *Cladocopium proliferum* (Butler et al. 2023) and *Durusdinium cf. trenchii* (LaJeunesse et al 2018) at densities of 3000 and 5000 cells mL^-1^ respectively. Illumination over each tank was provided by a single LED panel (Hydra® 52, Aqua Illumination®, USA). Upon settlement, a photon irradiance (400 – 700 nm) of 15 µmol photons m^-2^ s^-1^ was provided for the first 14 days and slowly increased to 150 µmol photons m^-2^ s^-1^ over three months. Recruits were fed once daily with enriched rotifers at a final density of 0.5 rotifers mL^-1^, and after one month, and instar I *Artemia salina* was also provided once daily at a final density of 0.5 nauplii mL^-1^.
