## Supplementary Table 2 for "Controlled “out-of-season” spawning of reef-forming corals in aquaria using offset environmental profiles"

**Table S2**: Fertilisation percentages for larvae cultures established from F1 coral colonies over spawning 2022 and 2023. Species key: *A. els* – *Acropora elseyi. A. hya* – *Acropora hyacinthus. A. lor* – *Acropora loripes. A. mil* – *Acropora millepora. M. aeq* – *Montipora aequituburculata.*

| **Year** | **Date** | **Species** | **Genotypes** | **Replicate 1** | | **Replicate 2** | | **Replicate 3** | | **Average fert. (%)** |
| --- | --- | --- | --- | --- | --- | --- | --- | --- | --- | --- |
|  |  |  |  | **Sample size** | **Embryos** | **Sample size** | **Embryos** | **Sample size** | **Embryos** |  |
| **2022** |  |  |  |  |  |  |  |  |  |  |
|  | 15/05/2022 | *A.mil* | 2 | 25 | 21 | 21 | 16 | 23 | 18 | 79.7 |
|  | 16/05/2022 | *A.els* | 3 | 50 | 17 | 57 | 18 | 58 | 18 | 32.1 |
|  | 10/06/2022 | *A.lor* | 3 | 23 | 13 | 28 | 19 | 87 | 58 | 65.2 |
|  | 13/06/2022 | *A.lor* | 9 | 121 | 91 | 43 | 34 | 60 | 47 | 76.8 |
| **2023** |  |  |  |  |  |  |  |  |  |  |
|  | 3/05/2023 | *M. aeq* | 3 | 46 | 4 | 52 | 5 | 51 | 7 | 10.7 |
|  | 4/05/2023 | *M. aeq* | 6 | 108 | 96 | 146 | 133 | 78 | 72 | 90.7 |
|  | 1/06/2023 | *M. aeq* | 3 | 104 | 98 | 81 | 77 | 67 | 61 | 93.7 |
|  | 1/06/2023 | *A.mil* | 2 | 89 | 67 | 41 | 32 | 53 | 43 | 77.6 |
|  | 1/06/2023 | *A.els* | 3 | 40 | 31 | 50 | 34 | 108 | 85 | 75.8 |
|  | 1/06/2023 | *A.lor* | 9 | 39 | 33 | 45 | 40 | 112 | 92 | 84.2 |
|  | 1/06/2023 | *A. hya* | 3 | 80 | 65 | 70 | 56 | 114 | 98 | 83.0 |
|  | 8/06/2023 | *A.lor* | 17 | 70 | 66 | 61 | 56 | 66 | 59 | 91.9 |
